# CRISPR disruption and UK Biobank analysis of a highly conserved polymorphic enhancer suggests a role in male anxiety and ethanol intake

**DOI:** 10.1101/572065

**Authors:** Andrew R. McEwan, Connor Davidson, Elizabeth Hay, Yvonne Turnbull, Johanna Celene Erickson, Pietro Marini, Dana Wilson, Andrew M. McIntosh, Mark J. Adams, Chris Murgatroyd, Perry Barrett, Mirela Delibegovic, Toni-Kim Clarke, Alasdair MacKenzie

## Abstract

Excessive alcohol intake is associated with 5.9% of global deaths. However, this figure is especially acute in men such that 7.6% of deaths can be attributed to alcohol intake. Previous studies identified a significant interaction between genotypes of the galanin (*GAL)* gene with anxiety and alcohol abuse in different male populations but were unable to define a mechanism. To address these issues the current study analysed the human UK Biobank cohort and identified a significant interaction (n=115,865; *p*=0.0007) between allelic variation (GG or CA genotypes) in the highly conserved human GAL5.1 enhancer, alcohol intake (AUDIT questionnaire scores) and anxiety in men that was consistent with these previous studies. Critically, disruption of GAL5.1 in mice using CRISPR genome editing significantly reduced *GAL* expression in the amygdala and hypothalamus whilst producing a corresponding reduction in ethanol intake in KO mice. Intriguingly, we also found evidence of reduced anxiety-like behaviour in male GAL5.1KO animals mirroring that seen in humans. Using bioinformatic analysis and co-transfection studies we further identified the EGR1 transcription factor, that is co expressed with *GAL* in amygdala and hypothalamus, as being important in the protein kinase C (PKC) supported activity of the GG genotype of GAL5.1 but less so in the CA genotype. Our unique study uses a novel combination of human association analysis, CRISPR genome editing in mice, animal behavioural analysis and cell culture studies to identify a highly conserved regulatory mechanism linking anxiety and alcohol intake that might contribute to increased susceptibility to anxiety and alcohol abuse in men.

## Background

The relationship between alcohol abuse and anxiety has been extensively explored ^1^ but the genomic mechanisms linking them remain poorly understood. Previous studies have shown that the galanin neuropeptide (encoded by the *GAL* gene) influences alcohol intake ^2, 3^ and anxiety-like behaviour^4^. Moreover, genetic analyses of different genotypes within the *GAL* locus succeeded in identifying an association with excess alcohol intake that was influenced by sex and anxiety^5^. Unfortunately, these studies were unable to define a mechanism to explain these interactions as the *GAL* coding region lacks non-synonymous polymorphisms^5^. Although there is a possibility that mis-regulation of *GAL* may affect alcohol intake and anxiety, little is known of the genomic mechanisms that modulate the expression of the *GAL* gene in the brain.

Based on the hypothesis that regions of the genome essential to species fitness are conserved through evolution ^6, 7^, we undertook a comparative genomic analysis of the genome surrounding the human *GAL* locus and succeeded in identifying a highly conserved enhancer sequence (hGAL5.1) 42 kilobases (kb) from the *GAL* gene transcriptional start site^8^. We demonstrated that GAL5.1 was active in galanin expressing cells of the hypothalamus including the PVN and dorsomedial hypothalamus (DMH)^8^. We also found that GAL5.1 contained two polymorphisms in perfect linkage disequilibrium (LD; rs2513280 (C/G) and rs2513281 (A/G)) that produced two different genotypes (GG and CA) within the human population and reported that the major GG genotype was significantly more active in primary hypothalamic neurones than the minor CA genotype ^8^. Analysis of these genotypes in human populations detected an association of the GG genotype and volume of alcohol intake in women in a small US cohort (n=138) ^9^. However, a second UK (n=2731) and US cohort (n=4064) based study, although initially identifying an association to increased frequency of binge drinking in teenagers, failed to reach significance after correction for multiple comparison ^10^. Because of the comparatively small size of these studies we interrogated a much larger human cohort (UK biobank) to better determine the association of specific allelic variants in GAL5.1 with levels of alcohol intake and anxiety in UK men and women. In addition, we used CRISPR genome editing to disrupt GAL5.1 in mice and examined the effects on the expression of flanking genes in areas of the brain including the hypothalamus and the amygdala to functionally link GAL5.1 activity to the expression of five flanking genes including *Gal.* Our study also characterises the effects of GAL5.1 disruption on ethanol consumption and anxiety-like behaviour in mice. A combination of bioinformatics (DNAseI hypersensitivity) and co-transfection studies also identified a transcription factor-DNA interaction underpinning GAL5.1 function and its response to signal transduction cues. This study highlights a mechanistic link between ethanol consumption and anxiety centred on the GAL5.1 enhancer that may contribute to the development of alcohol abuse and anxiety in men^11, 12^.

## Methods

### Genetic Association Study in humans

Genetic association analyses with alcohol consumption (units consumed per week) were performed using BGENIE, version 1 ^13^ with alcohol consumption as the outcome variable and age, sex, genotyping array and the first 20 genetic principal components as covariates. We then split the sample into males and females and repeated the association analysis, removing sex as a covariate. The mean alcohol intake in the total population was 15.3 units per week (S.D=16.0). The mean age of the population was 56.2 years (S.D. = 8.0). Men self-reported higher weekly alcohol intake compared to women (20.7 units (S.D.=18.4) vs 10.5 units (S.D.=11.5). Analysis of the association of alcohol intake and allelic variation at the rs2513280 locus was undertaken on a population of 345,140 individuals (UK Biobank (UKB); 183,921 females and 161,219 males all unrelated White British individuals).

To analyse the interaction effect of rs2513280 and anxiety on alcohol use behaviour in humans we also analysed a subset of the UK Biobank^14^ who had responded to a mental health questionnaire (MHQ) the results of which were made available to researchers in August 2017 ^15^. The MHQ had information on anxiety and in this follow-up alcohol use was ascertained using the AUDIT (Babor, 1991: The Alcohol Use Disorders Identification Test – Guidelines for Use in Primary Care. World Health Organ - Dep Ment Health Subst Depend 2001); a 10-item questionnaire used to measure both alcohol consumption and problematic drinking. AUDIT total scores range from 0-40 and the derivation of this measure in the UKB has been described previously in greater detail ^16^. For both sets of alcohol analyses, former and never drinkers were removed.

Anxiety was ascertained by asking participants “Over the last 2 weeks, how often have you been bothered by any of the following problems? Feeling nervous, anxious or on edge” [UKB data field 20506]. Anxiety was then dichotomized by comparing those answering ‘Not at all’ to individuals endorsing ‘Several days’, ‘More than half the days’ or ‘Every day’. After filtering the MHQ subset of individuals on those who were White, British and unrelated with and non-missing data there remained 115,865 individuals available for analysis. Linear regression models implemented in R were used to test the association between AUDIT score and rs2513280 genotype and the interaction effects of anxiety and sex. With AUDIT score as the outcome variable we fitted rs2513280, sex and anxiety as main effects and then a three-way interaction term of sex*anxiety*rs2513280. We also used twenty principal components as covariates to adjust for potential population stratification in the UK Biobank sample. rs2513280 genotype was modelled as a continuous variable (coded 0,1,2 corresponding to the number of C alleles carried).

### Generation of gRNA molecules by a novel annealed oligo template (AOT) method

Single guide RNA (sgRNA) molecules were designed to disrupt the GAL5.1 enhancer using the optimised CRISPR design tool (http://CRISPR.mit.edu/). sgRNA template was produced by annealing oligonucleotides to produce two different DNA templates as previously described ^17^ that included a T7 polymerase binding site and predicted guide sequence target sites spanning the most conserved central region of the mGAL5.1 enhancer (5’sgRNA; CTC CCT GGA GCA ATA TGA AG and 3’sgRNA; CCC GCT TTC ATG GCT CCC AA). These oligonucleotides were annealed and amplified using PCR to produce a 122 bp double strand sgRNA template. 100 ng of this template was used to produce sgRNA using a mMESSAGE mMACHINE T7 *in-vitro* transcription kit (Ambion) described in the manufacturer’s instructions and purified using a Megaclear kit (Ambion) with modifications as previously described ^18^.

### Production of genome edited mice

sgRNA molecules were microinjected at a concentration 10 ng/*µ*l each into the cytoplasm of 1-cell C57/BL6 embryos (Charles River) as described ^18^ together with 10 ng/*µ*l CAS9 mRNA (Life Technologies). Two-cell embryos were introduced into host CD1 mothers using oviduct transfer as previously described ^19^ and correctly targeted offspring were determined by PCR of earclip DNA using the following flanking primers (mGAL5for; AGTTAGGGCGCACACATCAA, mGAL5rev; CCGTGACTAACG GCTAATGC). These PCR products were purified and sequenced. Sequencing data was then analysed using the Blat tool (https://www.ncbi.nlm.nih.gov/pmc/articles/PMC187518/ “BLAST-like alignment tool”) of the UCSC browser to compare the sequence of the PCR product against that of the mouse genome (Fig 1F and G).

**Figure 1.**
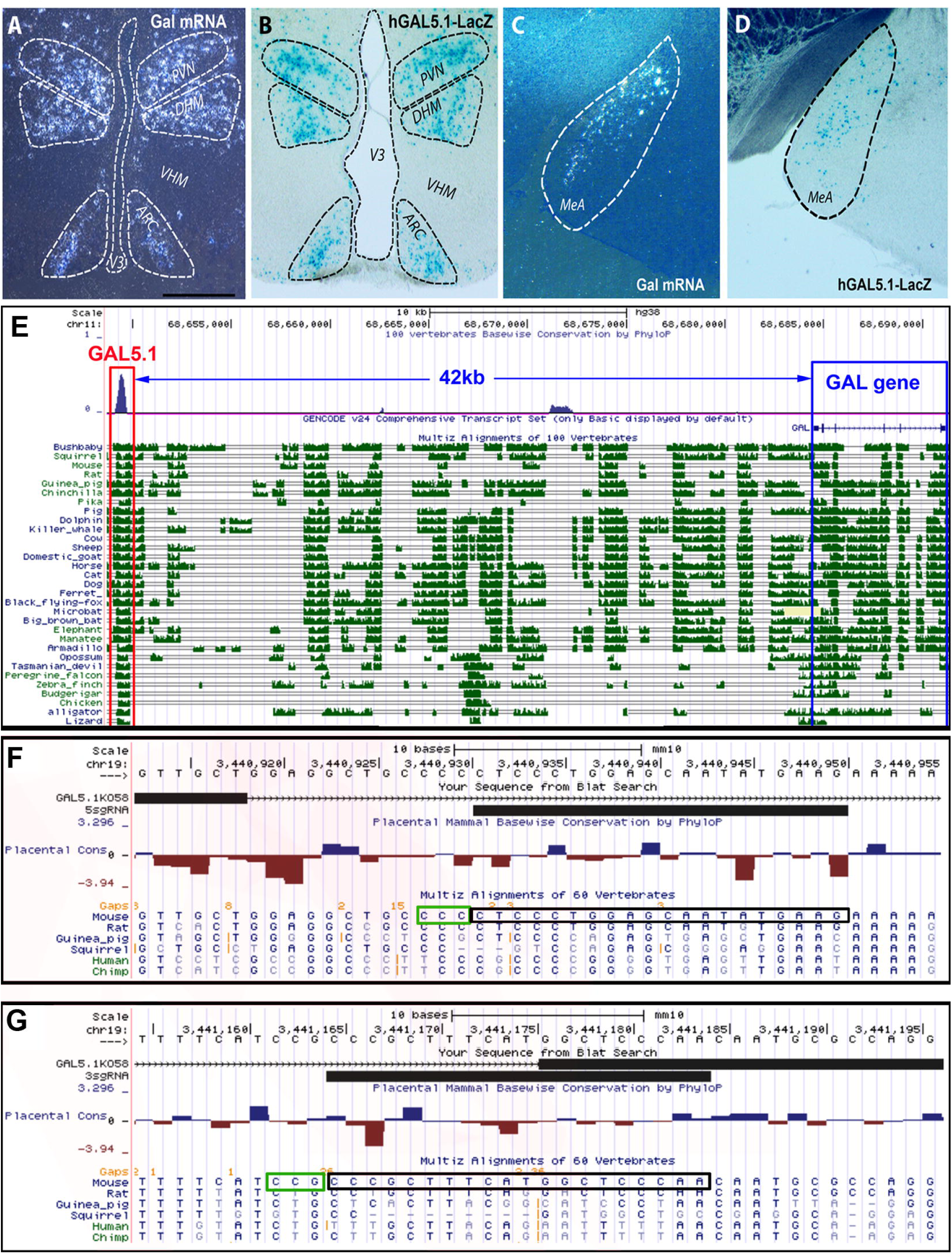
**A and C**, Dark field images of 10µm sections through **(A)** the hypothalamic region and **(C)** the amygdala region of wildtype mouse probed with S^35^ labelled *Gal* antisense RNA. **(B** and **D)** vibratome sections through **(B)** the hypothalamic and **(D)** the amygdala region of a mouse transgenic for the human GAL5.1-LacZ reporter construct (hGAL5.1-LacZ) demonstrating LacZ expression in cells of the periventricular (PVN) Dorsomedial (DMH) and arcuate (ARC) nuclei (V3, 3^rd^ ventricle) of the hypothalamus **(A and B)** and the (MeA) medial amygdaloid nucleus of the amygdala **(C and D). E.** Comparative genomic analysis of sequence flanking the human *GAL* locus demonstrating the position (numbered black scale bar at top), distance (blue arrows) and depth of conservation (green peaks) of the human GAL5.1 locus (red box) relative to the *GAL* gene (blue box). **F and G.** Sequence data derived from PCR products of ear clip DNA derived from a homozygote GAL5.1 KO mouse (thick black bar labelled GAL5.1KO58) blasted against the mouse genome (UCSC browser). The sequence disruption produced by introduction of CAS9/sgRNA is displayed relative to the PAM sequence (green box) of each sgRNA demonstrates the accuracy of **(F)** 5sgRNA and (**G)** 3sgRNA relative to the missing regions of the genome (thin black lines and chevrons). Electrophoresis gels demonstrating the difference in PCR product sizes are published elsewhere (Hay *et al.*, 2018).

### *In-situ* Hybridisation

Radioactive *in-situ* hybridisation was carried out on 10μm brain sections derived from wild type mice using radiolabelled *Gal* or Early Growth Response gene 1 (*EGR1 or Zif268))* antisense RNA probes as previously described ^20^.

### Quantitative reverse transcriptase-PCR

Brain tissues (Hypothalamus, amygdala, hippocampus and cortex) were dissected out of the whole brain and snap frozen on dry ice. Total RNA was extracted using the isolate II RNA minikit (Bioline). Quantitative reverse transcriptase-PCR (QrtPCR) to determine mRNA expression levels of genes flanking mGAL5.1 (Low-density lipoprotein receptor-related protein 5 (*Lrp5)*, Protein Phosphatase 6 Regulatory Subunit 3 (*Ppp6r3), Gal*, Tesmin *(Mlt5)* and carnitine palmitoyltransferase 1A *(Cpt1a*) was undertaken using gene specific primers (See table 1 in supplementary data) on derived cDNA using mouse Qrt-PCR primers as previously described ^21, 22^ using a Roche Light Cycler 480 with Roche SYBR green. All QrtPCR analyses were normalised using mouse primers specific to the Non-POU Domain Containing Octamer Binding protein (*Nono)* housekeeping gene that gave the most stable expression in all neuronal tissues analysed compared to βactin, Hypoxanthine-guanine phosphoribosyltransferase (HPRT) or Glyceraldehyde-3-Phosphate Dehydrogenase (GAPDH) genes.

**Table 1.** Interactions between different variables after analysis of the UK Biobank demonstrating a stronger than expected interaction (*p*=0.0008) between anxiety, sex, allelic variation at the rs2513280 locus and alcohol intake as measured by AUDIT score. **B.**

| Variable | Estimate | S.E. | P-value |
| --- | --- | --- | --- |
| Sex (male) | 0.34 | 0.005 | $<2 \times 10^{-16}$ |
| rs2513280 | 0.003 | 0.006 | 0.58 |
| Anxiety | 0.03 | 0.006 | $<3.7 \times 10^{-6}$ |
| Sex(male)*anxiety | -0.005 | 0.01 | 0.60 |
| Sex(male) *rs2513280 | -0.01 | 0.009 | 0.12 |
| Anxiety*rs2513280 | -0.02 | 0.01 | 0.10 |
| Sex(male)*anxiety*rs2513280 | 0.06 | 0.02 | **0.0007** |

### Animal studies

All animal studies were performed in accordance with UK Home Office guidelines. All mice used were sex and age matched (12-18 weeks) littermates. Phenotypic analysis of the morphology, general health, motility, behaviour and breeding success of these animals was also carried out. All animals were assigned random numbers after genotyping so that none of the operators knew the genotypes of the animals prior to testing.

### Alcohol preference studies

Preference for ethanol was tested by allowing animals free access to a choice of 10% v/v ethanol or pure water intake using a two-bottle choice protocol as described previously ^23^. Mice were housed in TSE home cage systems (TSE) that record liquid intake automatically. Initially mice were group housed (∼4 / cage) and habituated for 5 days to allow for adaptation to the monitored bottles. The mice were subsequently singly housed and habituated for a further 5 days prior to introduction of a second bottle containing the ethanol solution. Intake of both water and 10% ethanol solution were monitored at 30 min intervals over the course of the trial and recorded by TSE PhenoMaster Software (TSE-systems). Bottles were regularly swapped to reduce positional effects on consumption. A taste preference test to identify differences in sweet and bitter taste preferences between wild type and GAL5.1KO animals was performed by offering animals a choice of water, water and saccharine (2mM) or water and quinine (0.3mM) ^24^.

### Open Field test (OFT)

The OFT consisted of a square 30□cm (30□cm high) PVC open field arena positioned on a white base with overhead lighting applying 100 lx at the base^25^. Animals were transported individually to the testing room, habituated (30 mins) and released into the corner of the arena. Ambulatory activity was recorded in the open field for 300 s using an overhead video camera and the ANY-maze tracking software (Stoelting Europe). The software was used to define two zones in the open field at approximately 35 cm^2^ centre zone and 30 cm^2^ peripheral zone. Distance travelled, mean speed, Peripheral zone and no. centre entries, were determined by the software.

### Elevated zero maze (EZM)

The EZM consists of an annular dark-gray platform (60 cm in diameter) constructed of opaque Perspex divided into four equal quadrants. Two opposite quadrants were “open”; the remaining two “closed” quadrants were surrounded by 16 cm high dark, opaque black walls. Quadrant lanes were 5 cm in width. Overhead lighting applying 100 lx at the level of the maze. The movement of animals was tracked using a camera and ANY-maze tracking software. Distance travelled, average speed, number of crossings between light/dark zones and freezing episodes, freezing time and freezing latency were recorded.

### Novel suppressed feeding test (NSFT)

The NSFT assesses levels of anxiety by measuring the latency of a fasted animal to approach and eat a familiar food in an unfamiliar environment. The NSFT resembles the OFT except that animals are fasted for 16 hours before the test and the animal is then placed in the open field apparatus with a pellet of food fixed to a disc of filter paper and placed in the centre of the cage. The latency period between placing the animal in the test and first bite of the food pellet was recorded. The numbers of lines crossing and the speed and distance covered by the animal during the test was also recorded.

### GAL5.1 Deletion and expression constructs

Primers were designed that flanked the conserved putative EGR1 binding site within the GAL5.1 enhancer (ARM212-D2F; ATA GAT TTC AGA AAA GAA AGC TT, ARM213-D2R; TAA AAT GAC TGG CAT TAG AGC TC −3’). These were used in a PCR reaction with Q5 Hi-Fidelity polymerase (NEB, USA) in combination with the pLuc-GAL5.1 reporter construct previously described ^8^ as template to produce pLuc-GALΔEGR. The PCR product was ligated with T4 ligase prior to transformation into competent cells (Stratagene). The plasmid was sequenced to validate the removal of the putative EGR1 binding site. Expression constructs (pcDNA-EGR1 and pcDNA-PKCε) were obtained from Addgene.

### Cell culture and transfection studies

SH-SY5Y cells (94030304, ECAC) were cultured in Dulbecco’s Modified Eagle Medium (DMEM; Gibco,) containing low Glucose (5.5mM), L-Glutamine (4mM), and Sodium Pyruvate (1mM). Medium was previously supplemented with 10% (v/v) heat-inactivated Foetal Bovine Serum (FBS; Gibco) and 1% (v/v) Penicillin-Streptomycin (Pen-Strep; Gibco, UK). Cells were transfected with luciferase reporter plasmids DNA described above using jetPRIME as per manufacturer’s instruction (Polyplus Illkirch). Assays were normalised by co-tranfected with renilla luciferase plasmid pGL4.74 (Promega) or by assaying total protein in extract. After transfection, cells were treated with phorbol 12**-**myristate 13-acetate (PMA; 100nM in DMSO) or DMSO or PKC antagonist (GF10930X; Tocris) for 24 hours following which cells were lysed for dual luciferase analysis as per manufacturer’s instructions (Promega UK).

### Data analysis

From *in-vivo* pilot studies we calculated that a minimum of 6-12 animals per group would enable detection of a 25% difference between different parameters (ethanol intake, fat deposition) with 80% power using one-way ANOVA and/or general linear modelling. Statistical significance of data sets was analysed using either one-way analysis of variance (ANOVA) analysis with Bonferroni post hoc tests or using two tailed unpaired parametric Student *t*-test as indicated using GraphPad PRISM version 5.02 (GraphPad Software, La Jolla, CA, USA).

## Results

### The G-allele of rs2513280 associates with increased alcohol consumption in women

In order to test the validity of two previous association analyses^9, 10^ we investigated whether there was an association between allelic variants of the GAL5.1 enhancer and increased alcohol intake in a much larger cohort comprising 345,140 individuals (UK Biobank; 183,921 females and 161,219 males). In the total sample a marginal association between the G allele of rs2513280 (a proxy for both loci) and increased weekly alcohol intake in women was identified (b=0.008, s.e.=0.004, p=0.0463). When analysing males and females separately, the association between male alcohol intake and rs2513280 became non-significant (b=0.004, s.e.=0.006, p=0.478); however, in females the association was stronger and remained significant (b=0.012, s.e.=0.006, p=0.037).

Because of the known link between anxiety and alcohol abuse in men^26^ we further investigated the interaction of these variables with respect to the rs2513280 using a subset of the UK Biobank that also contained information relating to anxiety^15^. DNA was derived from blood and there are 64,465 females and 51,539 males in the anxiety cohort. There was an effect of male sex on AUDIT score with males having significantly higher AUDIT scores than females (**Table 1**, b=0.34, S.E.=0.005, p < 2 × 10^−16^). Individuals with higher levels of self-reported anxiety also had significantly higher AUDIT scores although there was no main effect of the rs2513280 polymorphism **(Table 1)**. No significant interactions were detected between sex and rs2153280 or anxiety and rs2513280 (**Table 1**). However, anxious males reported significantly higher AUDIT scores (*p*=0.0007) if they carried the major allele at the rs2153280 locus (**Table 1 and supplementary Fig 1**). Comparisons of analyses carried out using AUDIT-T, AUDIT-C and AUDIT-P are also shown in for comparison (supplementary data figure 2.)

### The GAL5.1 enhancer supports reporter gene expression in specific regions of the brain including the hypothalamus and amygdala

Cloning of the human GAL5.1 enhancer next to a LacZ reporter (hGAL5.1-LacZ) and production of transgenic reporter mice as previously described^8^ demonstrated expression of the LacZ reporter in cells of the periventricular (PVN), dorsomedial (DMH) and arcuate nuclei (ARC; **Fig 1B**) of the hypothalamus as well as the medial nucleus of the amygdala (MeA; **Fig 1C**) that corresponded to the expression patterns of the endogenous wild type mouse *Gal* gene both at a tissue-specific (**Fig 1A and C**) and cellular level ^8^.

### Rapid and accurate disruption of mGAL5.1 in mice was achieved using CRISPR/CAS9

To determine a functional role for GAL5.1 *in-vivo* we chose two target sequences (5’sgRNA and 3’sgRNA) that flanked the most highly conserved region of the mouse GAL5.1 homolog (mGAL5.1; chr19:3440931-3441185; **Fig 1E**) and used our previously described AOT method to produce sgRNAs that, when injected into mouse embryos, should induce a disruptive 230 bp deletion in mGAL5.1^17^. We microinjected AOT derived sgRNA and CAS9mRNA into the cytoplasm of 90 1-cell C57/BL6 mouse embryos. 90% of these embryos survived and were oviduct transferred into host female CD1 mice to generate a homozygous female and two heterozygous male mice that contained identical deletions within GAL5.1 following analysis by PCR and electrophoresis (mGAL5.1KO; **Fig 1F and G**) without any evidence of off-target effects ^27^. These mice were outbred on wildtype C57/BL6 mice to produce a colony of male and female mGAL5.1KO heterozygote mice that were subsequently used to produce the age/sex and homozygote wildtype and mGAL5.1KO littermate mice (determined by PCR of earclip DNA from each animal prior to testing) used in the current study. We also carried out an extensive phenotypical analysis of GAL5.1KO animals based on previous criteria ^28^ and found no significant observable phenotypic or health changes as a result of the genetic modification and GAL5.1KO mice were healthy and viable (see supplementary data 3 and 4).

### Disruption of mGAL5.1 reduces expression of *GAL* in amygdala, hypothalamus and hippocampus but does not affect the expression of flanking genes in hypothalamus

We compared the expression of the *Gal* gene in mRNA derived from various regions of the brain in wild type and mGAL5.1KO mice. Expression of *Gal* mRNA was strongest in the hypothalamus and the amygdala with some evidence of *Gal* mRNA expression in hippocampus and cortex (**Fig 2A and B**). We observed that expression of *Gal* became almost undetectable in all these tissues in mGAL5.1KO animals (**Fig 2A and B)** suggesting that mGAL5.1 is essential for a large proportion of *Gal* expression in these tissues.

**Figure 2.**
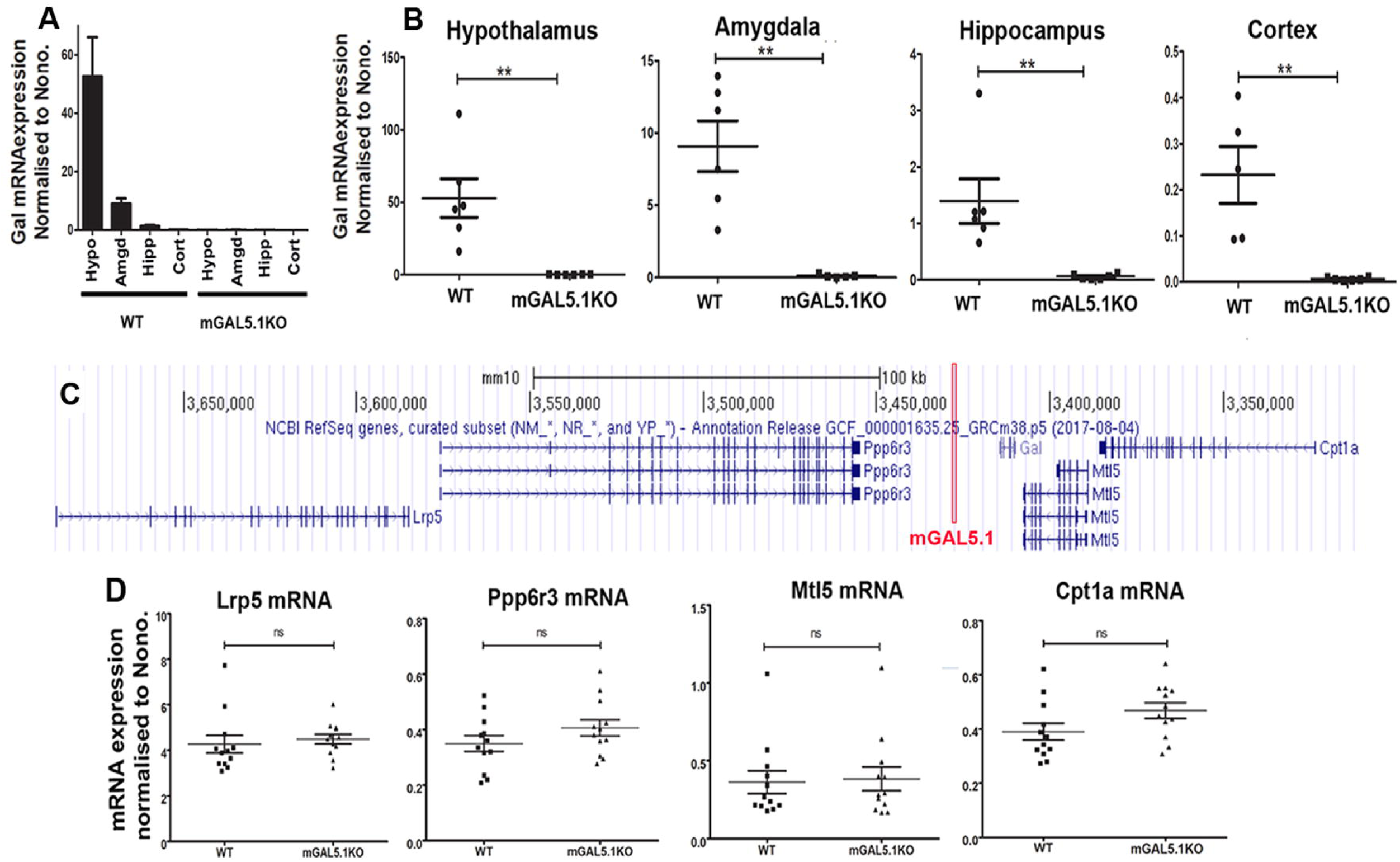
A. GAL5.1 specifically regulates expression of the *Gal* gene in the hypothalamus and amygdala but not flanking genes. **A.** Bar graph showing relative levels of *Gal* expression as measured by QrtPCR in different brain areas (Hypo, hypothalamus; Amyg, amygdala; Hipp, hippocampus, Cort; cortex) in wild type (WT) and mGAL5.1KO animals. **B.** Scatterplots demonstrating QrtPCR analysis of *Ga*l mRNA expression in RNA derived from hypothalamus, amygdala, hippocampus and cortex on total RNA derived from wild type (WT) or GAL5.1 knockout (mGAL5.1KO) animals(**;p<0.01). **C.** Scale diagram (UCSC genome browser) representing genes surrounding the mGAL5.1 enhancer (red box). Exons are displayed as thick blue lines and introns by thin blue lines punctuated by chevrons denoting the direction of transcription. Genomic coordinates on mouse chromosome 19 are highlighted by black perpendicular lines. **D.** Scatterplots showing QrtPCR analysis of cDNA derived from total hypothalamic RNA comparing mRNA expression of *Lrp5, Ppp6r3, Gal, Mtl5* and *Cpt1A* in wild type and GAL5.1KO animals normalised against the expression of the *Nono* gene (y-axis). (****, p<0.001, n.s., no significance).

We also analysed the influence of mGAL5.1 disruption on the expression of *Gal* and four other genes flanking the *Gal* locus namely *Lrp5, Ppp6r3* (5’ of mGAL5.1) *Mtl5* and *Cpt1a* (3’of mGAL5.1) that are all maintained in the same synteny block^29^ in both humans and mice spanning 536kb in humans and 379kb in mice (**Fig 2C**). Although we found strong down regulation of the *Gal* gene we saw no significant change of expression in any of the flanking genes suggesting a lack of requirement for mGAL5.1 for expression of these genes in the hypothalamus (**Fig 2D**)^30^.

### mGAL5.1KO animals exhibit a decreased preference for ethanol

Because the *GAL* gene has been shown to control intake of alcohol^2, 3^ and that a polymorphism within the GAL5.1 enhancer was associated with increased alcohol intake in women and in men who also reporting anxiety we tested the hypothesis that disruption of the mGAL5.1 enhancer would reduce preference for ethanol in mice. We provided ad-libitum access to bottles containing either water or water and 10% ethanol (both bottles sweetened with 0.05% saccharine) to singly housed mGAL5.1KO and WT male and female littermates mice and recorded intake for 10 days **(Fig 3A and B).** Both male and female mGAL5.1KO mice consumed significantly less 10% ethanol than wild type animals when given the choice (**Fig 3A**) and no significant difference in intake was detected between sexes. On average, wild type mice drank the equivalent of 3.7 grams of pure ethanol per kg per mouse per day whereas the GAL5.1KO mice only drank 1.1 grams (**Fig 3B**). To rule out the possibility that animals lacking the GAL5.1 enhancer had an altered preference for sweet or bitter tastes, that would skew their alcohol preference, we also carried out a taste preference test by offering wild type of GAL5.1KO animals a choice of water or water sweetened with saccharine or made bitter with quinine. We observed no evidence of changing preferences for bitter or sweet tastes as a result of GAL5.1 disruption (Supplementary figure 5).

**Figure 3.**
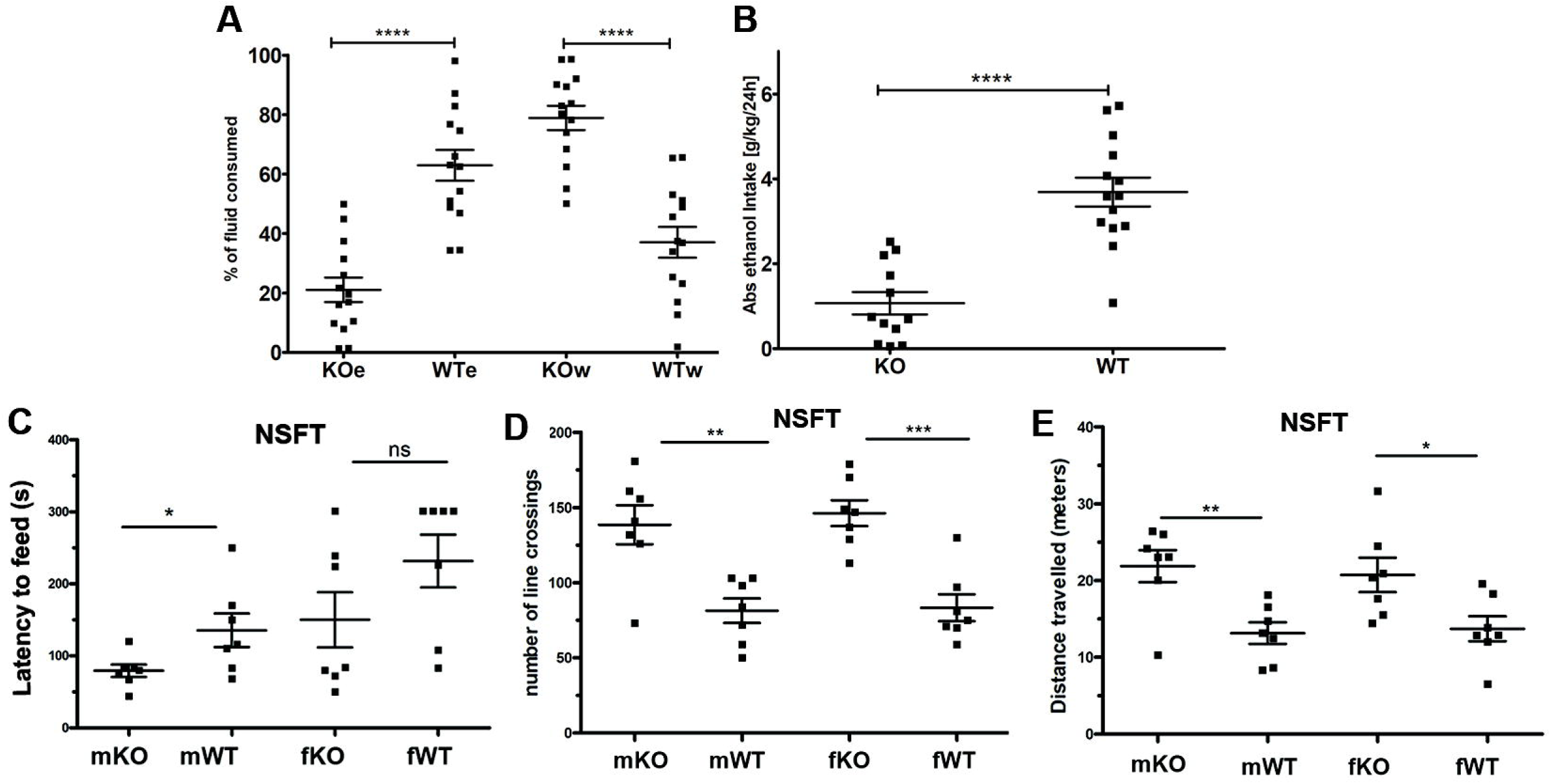
GAL5.1 disruption decreases ethanol intake and modulates anxiety-like behaviour in the marble burying test and the novelty suppressed feeding test. **A.** Scatterplot demonstrating the percentage intake of 10% ethanol in male and female mGAL5.1KO (KOe) and wild type litter mates (WTe) compared to water alone in mGAL5.1KO mice (KOw) and wild type littermates (WTw; error bars, SEM, n=12-14; ****, p<0.0001; F=30.83; d.f.=3) over the 10 days of the experiment. **B.** Scatterplot demonstrating ethanol intake in mGAL5.1KO (KO) male and female mice compared to wild type littermates (WT) expressed as grams of ethanol consumed per kilogram of mouse per day (error bars= SEM; n=12-14, ****, p<0.0001; t=5.918; d.f.=24). **C**. Scatter plots demonstrating the number of marbles buried in 30 minutes by male knockout (mKO), male wild type (mWT), female knockout (fKO) and female wild type (fWT) mice as analysed by the marble burying test (MBT; n=7-10; F=19.28; d.f.=3). **D-F**. Scatter plots demonstrating latency to feed (**D**; F=4.617; d.f.= 3), number of line crossings (**E**; F=12.46; d.f.=3) and distance travelled (**F**; F=5.965; d.f.=3) as analysed using the novelty suppressed feeding test (NSFT; n=7; error bars= SEM; n.s., not significant; *, p =0.05; **, p<0.01; ***, p<0.005; ****, p<0.0001).

### mGAL5.1KO mice exhibit an increase in marble burying and decreased latency to feed in the NSFT

Both the MBT and the NSFT have been widely used to assess levels of anxiety-like behaviour in mice. Using the MBT we observed that both male and female mGAL5.1KO animals buried significantly more marbles within the 30-minute period of the test than did their wild type littermates (**supplementary data 6**) consistent with reduced anxiety. In addition, we observed decreased latency in the time taken for fasted mGAL5.1KO animals to take their first bite of the food pellet within a novel environment although this only achieved significance in the mGAL5.1KO males (**Fig 3D**). Intriguingly, both males and female mGAL5.1KO mice demonstrated a significant increase in both numbers of line crossing between the peripheral zone and the centre zone (**Fig 3E**) as well as the distance travelled (**Fig 3F**) within the arena during the duration of the test.

### Male mGAL5.1KO mice demonstrate reduced anxiety-like behaviour in the OFT and the EZM tests

The OFT has been previously used as a test for anxiety-like behaviour ^31^. We first noticed a significant increase in the amount of centre entries displayed by male mGAL5.1KO mice compared to litter matched wild type animals (**Fig 4A**). We also noted an increase in the proportion of the distance moved by GAL5.1KO mice (**Fig 4B**) as well as the proportion of time spent within the centre zones (**Fig 4C**). No significant difference in these behaviours was observed between female WT(fWT) and mGAL5.1KO (fKO) mice exposed to the OFT (**Figs 4A-C**)

**Figure 4.**
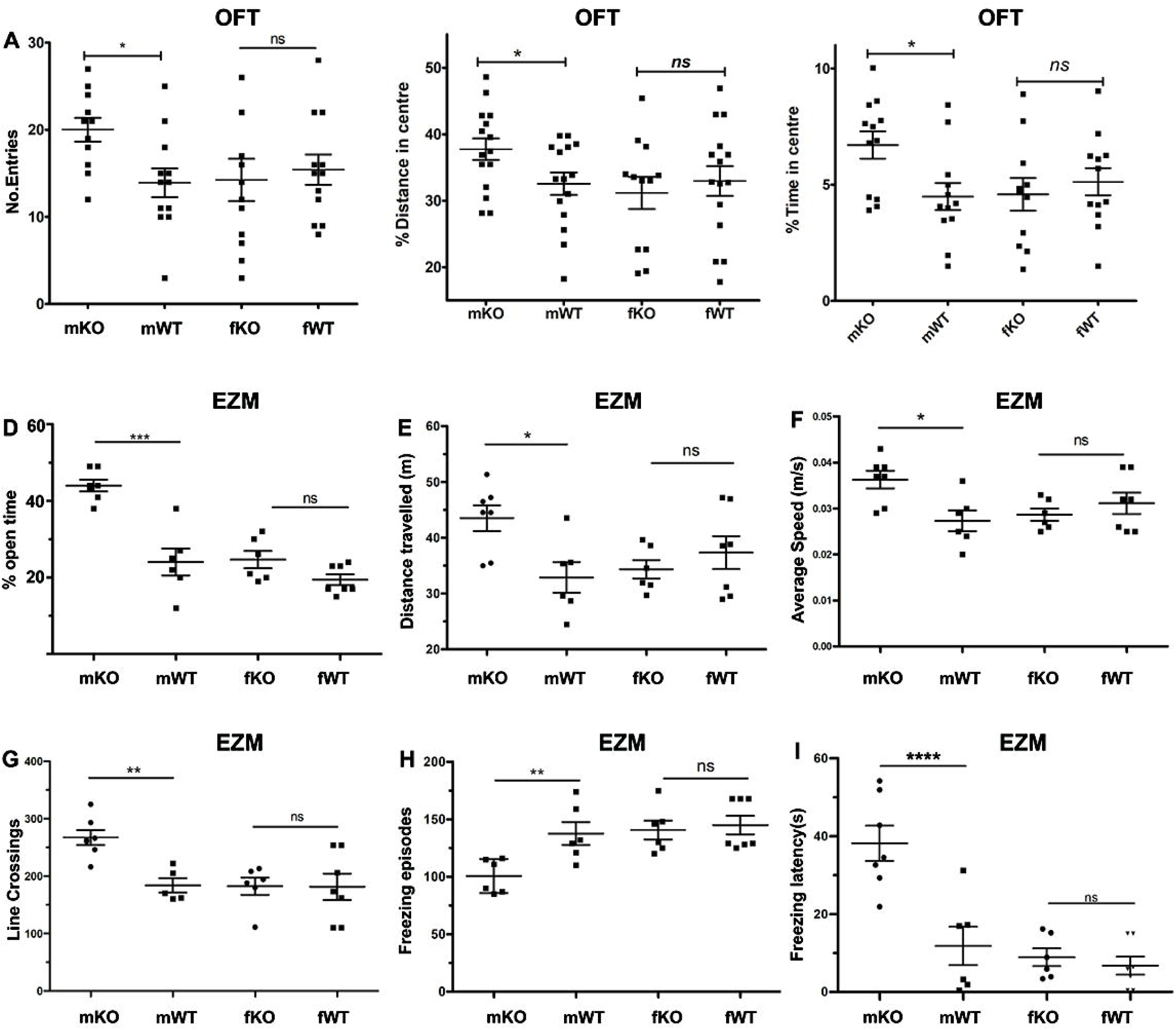
mGAL5.1 deletion affects sex specific aspects of anxiety-like behaviour in the open field test and the elevated zero maze. **A-C.** Scatterplots demonstrating increased number of centre entries (**A;** F=2.290, d.f.=3), percentage distance moved in the centre zone (**B;** F=2.583, d.f.=3) and the percentage of time in the centre zone (**C;** F=2.176, d.f.=3) of male GAL5.1KO mice (mKO) compared to male wild type mice (mWT) using the open field test (OFT). Female GAL5.1KO mice (fKO) and wild type mice (fWT) demonstrated no significant behavioural difference in the OFT(**A-C**). **D-I.** Scatterplots demonstrating changes in the time spent in the open quadrants of the elevated zero maze (EZM) (**D;** F=25.80, d.f.=3), the distance travelled (**E;** F=3.605, d.f.=3) the average speed (**F;** F=3.846, d.f.=3), the numbers of crossings between the light and dark zones (**G;** F=6.498, d.f.=3), the number of freezing episodes (**H;** F=6.088; d.f.=3) and the time until the first freezing event (**I;** f=16.48; d.f.=3) by mKO and mWT animals. Female GAL5.1KO mice (fKO) and wild type mice (fWT) demonstrated no significant behavioural difference (**D-I**) in the EZM. (OFT, n=10-12; EZM, n=7; n.s., not significant; *, p =0.05; **, p<0.01; ***, p<0.005; ****, p<0.0001).

The EZM measures the innate fear of small rodents for open areas against the security of a closed area and is a development of the elevated plus maze ^32^. We observed that male mice placed into the EZM displayed a significantly greater percentage of their time within the open quadrants of the maze (**Fig 4D**). In addition, male mice travelled a greater distance (**Fig 4E**) at a marginally faster speed (**Fig 4F**) whilst crossing between the light and dark sides with greater frequency (**Fig 4G**). Much of the difference in overall speed and distance between WT and KO males were attributable to the observed reduction in freezing behaviour (frequency and latency) displayed by male mGAL5.1KO animals compared to wild type animals (**Figs 4H and I**). No significant difference was observed between female mice exposed to the EZM (**Figs 4D-I**).

### EGR1 interaction and modulation of the PKC response varies with GAL5.1 genotype

To identify transcription factors that were involved in modulating GAL5.1 activity we undertook a bioinformatic analysis of the GAL5.1 enhancer using ENCODE data on the UCSC browser (**Fig 5A**). ENCODE identified highly conserved regions of GAL5.1 that were sensitive to DNAse1 digestion; a diagnostic of open, transcriptionally active chromatin, in several different cell lines **(Fig 5A and B).** This analysis also highlighted the presence of a highly conserved binding consensus of EGR transcription factors that lay within a DNAseI hypersensitive region (**Fig 5A and B**). Furthermore, EGR1 (AKA Zif268) is expressed in the PVN (**Fig 5C**) and is up-regulated in the ARC in response to anorexia stimulated melanocortin signalling^33^. We produced luciferase reporter constructs containing the GAL5.1(GG) enhancer and a derivative of GAL5.1(GG) lacking the conserved EGR binding site (pLuc-ΔEGR1). These were transfected into a neuroblastoma cell line (SH-SY5Y) in the presence of an empty expression vector (pcDNA3) or an expression vector expressing the EGR1 protein (pcDNA3-EGR1, Addgene). These experiments demonstrate that expression the EGR1 transcription factor in neuroblastoma cells significantly increased GAL5.1 activity (**Fig 5D**) and that the putative EGR1 binding site identified in GAL5.1 is critical to this interaction. In order to confirm that the effects of PMA on GAL5.1, as previously reported ^8^, were indeed PKC specific we tested the effects of different concentrations of the PKC antagonist GF10930X on the PMA driven up-regulation of the GAL5.1(GG) genotype. We showed that PMA induction was significantly reduced following co-treatment with 1000nM concentration of the PKC antagonist **(Fig 5E).** Further analysis comparing the effects of EGR1 expression on the GAL5.1(GG) and GAL5.1(CA) genotype reporter constructs showed that, whilst the GG genotype responded strongly to EGR1 expression in SH-SY5Y cells, the CA genotype did not (**Fig 5F**). In addition, the PMA induced response of the CA genotype was blunted in comparison to the GG genotype (**Fig 5F**).

**Figure 5.**
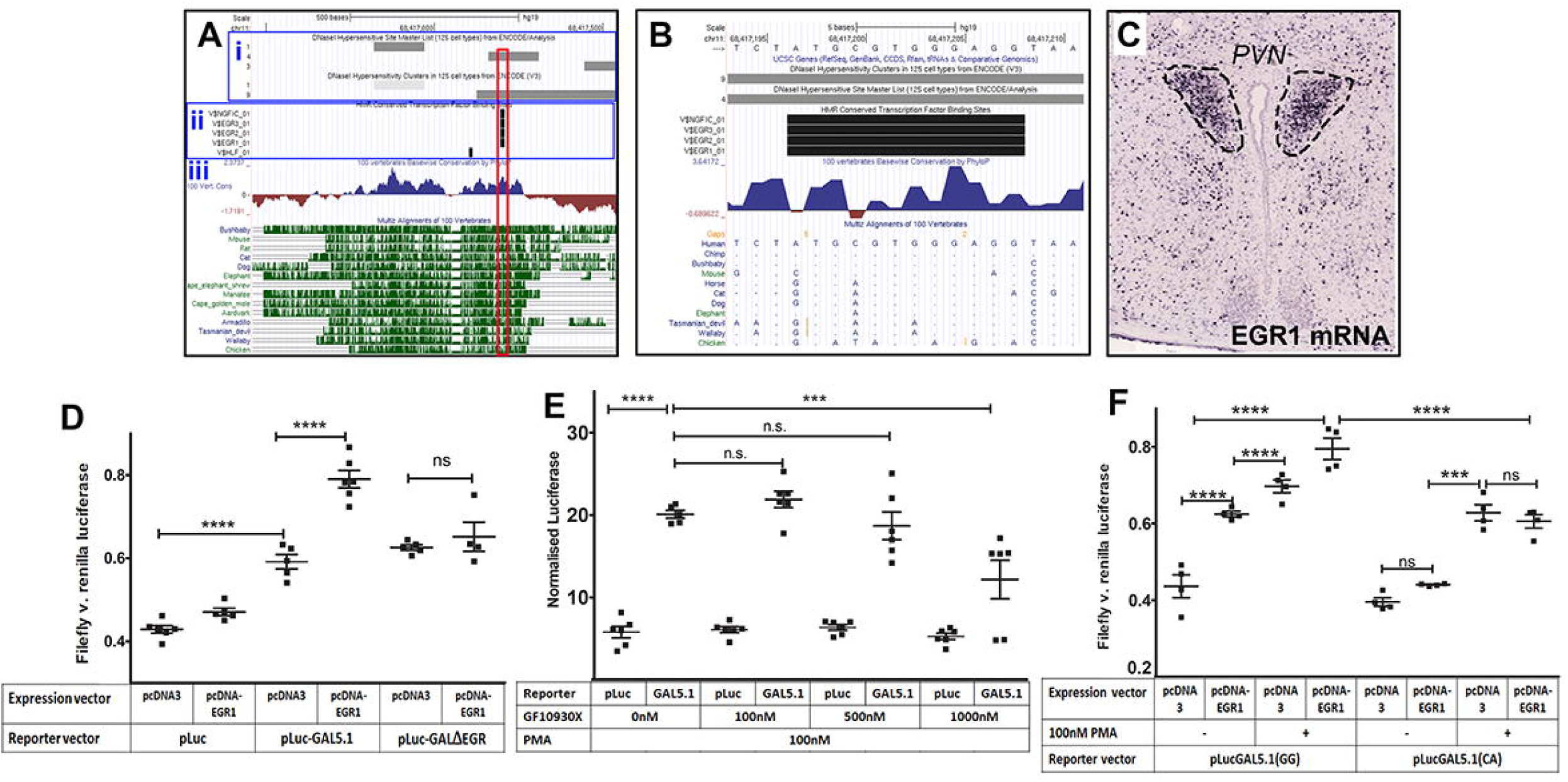
EGR1 interaction and modulation of the PKC response varies with GAL5.1 genotype. **A.** UCSC browser output of 800bp surrounding the GAL5.1 enhancer showing regions of DNAseI hypersensitivity (**Ai**; filled grey bars), conserved transcription factor binding consensus sequences (**Aii**; filled black boxes) and degree (**Aiii;** blue peaks) and depth (A**iii;** green lines) of sequence conservation. **B.** UCSC output showing hypersensitivity sites (grey bars), conserved EGR1 consensus sequences (black bars) and degrees of conservation (Blue peaks). Multiz alignment of conserved consensus sequences demonstrates degrees of conservation at the base pair level where dots represent identical base pair to human. **C.** *In-situ* hybridisation demonstrating the expression of *Egr1* mRNA within the periventricular nucleus (PVN) of the mouse hypothalamus. **D.** A scatterplot demonstrating dual luciferase data derived from SH-SY5Y cells transfected with different combinations of pcDNA3 (empty expression vector), pLuc (empty reporter vector), pcDNA-EGR1 (Expression vector expressing EGR1 transcription factor), pLuc-GAL5.1 (reporter vector containing the GAL5.1 enhancer), pLuc-GalΔEGR (reporter construct containing the GAL5.1 enhancer lacking the EGR binding consensus shown in **B** (n=5-6; F=60.47; d.f.=5). **E.** A scatterplot demonstrating the effects of different concentration of the PKC antagonist GF10930X on the PMA stimulated activity of the pLuc-GAL5.1 plasmid (GAL5.1) compared to the empty luciferase vector (pLuc; n=6, **, p<0.01; ***, p<0.005; F=37.62; d.f.=7). **F**. A Scatterplot comparing the effects of co-transfection of an EGR1 expressing plasmid (pcDNA-EGR1) and/or PMA treatment on cells co-transfected with a renilla luciferase expressing plasmid and a firefly luciferase reporter constructs containing either the GG or CA genotypes of the GAL5.1 enhancer (n=4, ns; no significance, **\*\*\***;p<0.005; **\*\*\*\***;p<0.001; F=54.91; d.f.=7).

## Discussion

Understanding the processes that modulate preference for ethanol is particularly pressing in men where 7.6% of all male deaths globally can be attributed to alcohol abuse through, accidental death, cardiovascular disease, liver damage and cancer ^34^. Anxiety remains one of the most frequent co-morbidities associated with excessive alcohol intake^35^ and identifying and understanding the genomic mechanisms that link anxiety with excessive alcohol intake will be an important component in understanding and combating alcohol abuse^26^. A previous genotype analysis of polymorphisms around the *GAL* gene in two different populations (Finns and Plains Native Americans) identified robust associations with alcohol intake and specific genotypes ^5^. Intriguingly, this study also uncovered evidence of a sex specific role for anxiety in modulating alcohol intake ^5^. Unfortunately, the authors were unable to identify a molecular mechanism that could account for their findings. The current study explored the problem from the initial standpoint of functionality. This was achieved by analysing the effects of polymorphisms known to change the activity of the highly conserved GAL5.1 enhancer sequence found 42kb from the *GAL* gene ^8^.

We began our analysis of a possible role for the GAL5.1 enhancer in alcohol intake by interrogating the UK Biobank cohort to determine the validity of previous conflicting smaller scale association studies ^9, 10^. Our analysis supported observations by Nikolova et al (2013) who reported an association between the G-allele of GAL5.1 and increased alcohol intake in women ^9^ but not in men. Because anxiety has been identified as an important variable influencing alcohol intake in humans^1^ and had also been identified as being an important variable in modulating the role of the *GAL* gene in alcohol intake^5^, we carried out a deeper analysis of the rs2513280 polymorphism UK biobank by stratifying alcohol intake with anxiety. We were interested to find a significant association between reported anxiety and drinking behaviour in men which was not observed in women. Although this observation was entirely consistent with previous reports of sexual dimorphism on the influence of the *GAL* locus on anxiety and alcohol intake^5^ it contrasted with an observed reduction of ethanol intake only in female GAL gene deletion mice ^36^. Whilst these results could be explained by the maintenance of the GAL gene deletion mouse line on a pure 129Ola/Hsd genetic background; a strain which displays a relatively low alcohol intake compared to the C57/BL6 line used in the current study ^37^, there is also a likelihood that, because disruption of the GAL5.1 enhancer does not remove all expression of the Gal gene, many of the sex specific differences observed may also be explained by the presence of residual sex-specific Gal gene expression in different parts of the brain.

To further explore the role of GAL5.1 in ethanol intake we disrupted GAL5.1 in mice using CRISPR/CAS9 genome editing. Using QrtPCR of different brain regions of GAL5.1 knock out mice we found that disruption of GAL5.1 virtually blocked expression of *Gal* mRNA expression in hypothalamus, amygdala, hippocampus and cortex consistent with a requirement for GAL5.1 for expression of *Gal* in these brain regions. Moreover, analysis of gene expression of four other flanking genes suggested that GAL5.1 is specific in its modulation of the expression of the *Gal* gene in the hypothalamus; although we are unable to rule out a possible role for GAL5.1 in regulating the expression of these genes in other tissues. However, the most striking outcome of the current study was the observation that CRISPR/CAS9 disruption of GAL5.1 significantly reduced preference for ethanol in mice demonstrating a key role for GAL5.1 in ethanol preference and intake.

Anxiety is one of the most frequently reported co-morbidities of alcohol abuse ^1, 35^. It is intriguing, therefore, that genetic analyses of different genotypes within the *GAL* locus succeeded in identifying an association with excess alcohol intake that was influenced by sex and anxiety^5^. These observations are supported by our current analysis of the UK biobank human cohort that also identified a significant association of a GAL5.1 enhancer polymorphism with alcohol abuse when stratified against sex and anxiety. We therefore asked whether our CRISPR/Cas9 knockout of GAL5.1 altered anxiety-like behaviour in mice and whether there was evidence of the apparent sexual dimorphisms observed in previous studies^5^ and in our UK Biobank analysis. Our initial analysis of mGAL5.1KO mice using the MBT demonstrated that removal of mGAL5.1 increased the tendency of both male and female mice lacking GAL5.1 to bury marbles; an accepted diagnostic of anxiety-like behaviour but also of obsessive compulsive disorder^38^. Interestingly, we also observed greatly increased numbers of line crossing as well as more distance travelled by both sexes of mGAL5.1 mice exposed the NSFT (AKA hyponeophagia) which suggests that GAL5.1 may play a role in exploratory foraging behaviour^39^. However, we only observed a decrease in feeding latency in the NSFT in males lacking GAL5.1 that first suggested the possibility that some aspects of anxiety-like behaviour in mice are modulated by GAL5.1 in a sex specific manner. This hypothesis was further reinforced by observations of the behaviours of mGAL5.1KO mice exposed to the OFT who spent proportionately more time in the centre of the OFT and proportionally moved a further distance. Further support for this observation came from the EZM whereby male mGAL5.1KO mice spent a significantly larger proportion of their time in the open quadrants of the EZM. Notably, these mice also displayed a reduced number of freezing episodes providing further evidence of a reduced fear response ^40^. The observed sex specific reductions in anxiety demonstrated by mGAL5.1KO mice in the NSFT, the OFT and the EZM are consistent with the sexual dimorphism observed in our stratified analysis of the UK biobank cohort and in previous analysis of polymorphisms around the *GAL* gene^5^. It is obvious that many more behavioural aspects of anxiety, such as social interaction, stress and conditioned fear response, need to be explored in these animals to better comprehend the specific role played by GAL5.1 in anxiety-related behaviour. Moreover, critics of the MBT and the NSFT correctly point out that the results of these tests may be skewed by an increase in compulsive behaviour and appetite respectively. When the results of all our tests are considered together, however, our study strongly suggests that the GAL5.1 enhancer plays an important role in modulating significant sex specific aspects of anxiety-like behaviour and ethanol intake in both mice and humans.

Even though we have identified phenotypic differences in the human population reflecting findings within our mouse models, we are also aware that the changes in our human cohort (2 pase pair change in hGAL5.1) are not strictly equivalent to those induced in our mouse models (230bp deletion in mGAL5.1) and we expected that our mouse deletion would produce a more extreme phenotype than that seen in humans. Indeed, we recognise that only by recreating both human the CA and GG genotypes in mice and comparing their phenotypic effects might a truly equivalent comparison be made. Nevertheless, our current study represents a major step towards defining a better understanding the role of GAL5.1 in mammalian ethanol intake and mood and defines a clearer path to identification and analysis of other conserved enhancers involved in modulating health and disease.

Closer analysis of the GAL5.1 sequence using a combination of bioinformatics and cell co-transfection studies suggested a functional interaction between the EGR1 transcription factor and a highly conserved EGR binding site within a highly conserved region of GAL5.1. EGR1 has a high affinity for DNA and is known to be able to bind DNA even when methylated 41. However, the literature describing the effects of ethanol on EGR1 expression and activity could be clearer. For example, it was reported that EGR1 binding to DNA was reduced following ethanol exposure^42^. In contrast, more recent reports suggest that expression of EGR1 in the brain was increased in animals exposed to ethanol ^43, 44^. From a mechanistic perspective it is possible that EGR1 acts as a “pioneer factor” by being one of the first transcription factors that bind to the closed and methylated GAL5.1 locus thus activating its tissue specific activity. Interestingly, we observed that the CA allele of GAL5.1 did not respond to EGR1 expression and its PKC response was blunted. This is an interesting observation as the EGR1 binding site is 100bp from the closest of the SNPs within GAL5.1 (rs2513281). We propose that the EGR1 protein forms part of a larger protein complex that interacts across the whole of the most conserved regions of the GAL5.1 enhancer. It is therefore possible that EGR1 binding and GAL5.1 function is dependent on binding of another, as yet unidentified, protein whose binding is interrupted by one of the allelic variants of GAL5.1 thus affecting EGR1 binding.

We also explored the possible interaction of PKC pathways in the activity of the GAL5.1 enhancer. Previous analysis demonstrated that the non-specific PKC agonist PMA increased the activity of GAL5.1 in primary hypothalamic cells^8^. In the current study we support these previous studies by showing that antagonism of PKC decreases the effects of PMA on GAL5.1 activity. We have also expanded our analysis to show that upregulation of PKA pathways increase the effects of EGR1 expression on the activity of the GG genotype of GAL5.1 but not in the less active and protective CA genotype; an observation that may allow us to design personalised therapies. Our best candidate for the PKC isoform responsible for modulating GAL5.1 activity, and to which we may target future drug therapies, is PKC epsilon (PKCε) which is expressed in the amygdala and PVN ^45, 46^. Moreover, genetic deletion of PKCε reduces ethanol intake ^45^ and anxiety like behaviour in mice ^46^. Future work will focus on determining the possible relevance of this isoform in the activity of the GAL5.1 enhancer and determining whether antagonism of this, and other PKC isoforms, may reduce the activity of the GG genotype to that of the protective CA genotype thus opening the possibility of developing a novel personalised anxiolytic therapy that reduces ethanol intake in men.

## Conclusion

To the best of our knowledge, this is the first time that the unique combination of techniques used in our study; human association analysis, comparative genomics and CRISPR/CAS9 genome editing, have been used to establish a functional role for a tissue-specific enhancer region in alcohol selection and mood in living animals. This study is given even greater impact by our analysis of the UK biobank cohort that demonstrates a link between increased alcohol intake and anxiety in males paralleling that seen in our CRISPR derived models. The current study also goes some way to addressing the age-old question of whether alcohol abuse causes anxiety (substance-induced anxiety model), or whether alcohol use is caused by anxiety (self-medication model), by providing supporting evidence that a common mechanism contributes to both behaviours (common-factor model) ^1, 35^.

Placing the current study within the wider context of understanding the mechanistic basis of complex human disease it is clear that an important step has been made in bridging the gap between association analyses and mechanism especially in light of the fact that the majority of associated SNPs generated by GWAS are in the non-coding genome^47^. Although histone markers (e.g. H3K4me1) and GWAS can identify candidate regulatory regions affected by disease associated polymorphic variation on a genome wide level the current study serves to emphasise that there is also merit in the use of comparative genomics and functional characterisation of the cell specific activity of putative regulatory elements using CRISPR genome editing in cell based and whole animal systems.

## Acknowledgements

We would like to thank Wenlong Huang, Giuseppe D’Agostino and the staff of the University of Aberdeen Medical Research Facility for their help and advice with the animal tests. AMcE was funded by BBSRC project grant (BB/N017544/1) and EH was funded by Medical Research Scotland (PhD-719-2013). PB and DW are funded by the Scottish Government Rural and Environment Science and Analytical Services Division to the Rowett Institute. Association studies were conducted using the UK Biobank Resource: application number 4844 and was supported by a Wellcome Trust Strategic Award ‘Stratifying Resilience and Depression Longitudinally’ (STRADL) (Reference 104036/Z/14/Z). Dedicated to the memory of my brother Angus MacKenzie (1963-2018).

