## Supplementary data 1 for "CRISPR disruption and UK Biobank analysis of a highly conserved polymorphic enhancer suggests a role in male anxiety and ethanol intake"

| Primer name | Sequence (5’-3’) | Product length | Mouse genome coordinates. |
| --- | --- | --- | --- |
| mPPP6R3_FOR | TTGCAGACCAAGACGACATTG | 236bp | chr19:3469738-3473793 |
| mPPP6R3_REV | TTCTTCACTGTCCGTACTGCC |  |  |
| mLRP5_FOR | AAGGGTGCTGTGTACTGGAC | 220bp | chr19:3659293-3659512 |
| mLRP5_REV | AGAAGAGAACCTTACGGGACG |  |  |
| mMTL5_FOR | CGGGATGAGTTGCCGGTTC | 208bp | chr19:3389073-3389280 |
| mMTL5_REV | CGGGAAGTAACGACGATAACAC |  |  |
| mGal_FOR | GGAAGTGTTGATGTGCCCCT | 253bp | chr19:3409992-3411558 |
| mGal_REV | GCAGAGAACAGACGATTGGC |  |  |
| mCPT1A_FOR | CTCCGCCTGAGCCATGAAG | 100bp | chr19:3349262-3352503 |
| mCPT1A_REV | CACCAGTGATGATGCCATTCT |  |  |
| mNoNo_FOR | GCCAGAATGAAGGCTTGACTAT | 105bp | chrX:101437372-101439585 |
| mNoNo_REV | TATCAGGGGGAAGATTGCCCA |  |  |

**Supplementary data 1.** Sequence, product length and mouse genomic coordinates of primers used in QrtPCR analysis of mouse hypothalamic mRNA. Primer sets were derived from validated primer sets archived on PrimerBank (https://pga.mgh.harvard.edu/primerbank/).
