## Supplementary data 2 for "CRISPR disruption and UK Biobank analysis of a highly conserved polymorphic enhancer suggests a role in male anxiety and ethanol intake"

| Association between rs2513280 and AUDIT scores | | | |
| --- | --- | --- | --- |
| Measure | Beta | SE | P |
| AUDIT-T | -0.0039 | 0.004 | 0.303 |
| AUDIT-C | -0.0029 | 0.003 | 0.397 |
| AUDIT-P | -0.0032 | 0.004 | 0.366 |
| Interaction analysis of rs2513280*anxiety*sex and AUDIT scores | | | |
| AUDIT-T | 0.057 | 0.016 | 0.0007 |
| AUDIT-C | 0.045 | 0.015 | 0.003 |
| AUDIT-P | 0.057 | 0.016 | 0.0004 |

**Supplementary data 2A.** An analysis of rs25132380 with the AUDIT subscales and presented for comparison to the full AUDIT score (AUDIT-T). We report no association between rs2513280 and any AUDIT subscale, reflecting the results reported for AUDIT-T in the current manuscript. The interaction between SNP*sex*anxiety is similar whether AUDIT-C or AUDIT-P is analysed suggesting these findings pertain to both consumption and problem drinking. We have added these to the supplementary materials but refrain from interpreting too far in the manuscript. AUDIT-C does reflect consumption but also includes a question on binge drinking and therefore also captures aspects of hazardous drinking. Whether these human findings are more relevant to consumption or dependence may be better explored in a clinically ascertained sample.

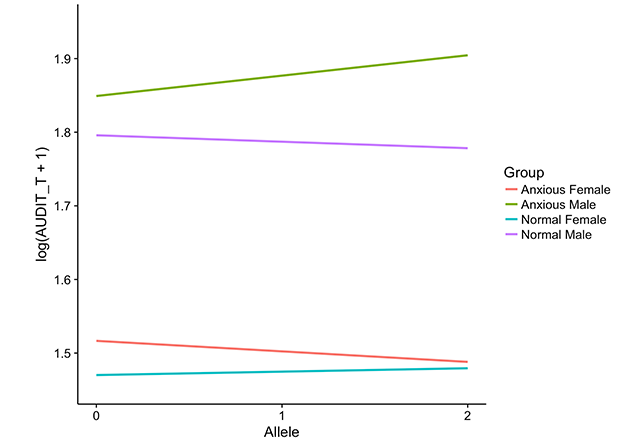

**Supplementary data 2B**. Graph showing the linear regression line for the 3-way interaction between sex, anxiety and rs2153280, with allele count representing the number of C alleles an individual carry and the y-axis showing the natural log transformed AUDIT-T score.
