## Supplementary data 3 for "CRISPR disruption and UK Biobank analysis of a highly conserved polymorphic enhancer suggests a role in male anxiety and ethanol intake"

**Supplementary data 3; Health screen for the ΔGal 5.1 (GAL5.1KO) mouse strain:**

| **Screening protocol** | **observation** | **Notes** |
| --- | --- | --- |
| Embryonic Lethal | No |  |
| Weaning behaviour | Normal |  |
| **Screening for visible birth defects** |  |  |
| Body size | Normal |  |
| Eye size | Normal |  |
| Eye colour | Normal |  |
| Skin colour | Normal |  |
| Skin texture | Normal |  |
| Stripes striations | No |  |
| General activity | Normal |  |
| Microagathia | No |  |
| agnathia | No |  |
| Short head | No |  |
| Scoliosis | No |  |
| Hare lip | No |  |
| Tail bend | No |  |
| Poly/syndactyly | No |  |
| Fused toes | No |  |
| Limb shape, length | No |  |
| Blebs or bruising | No |  |
| Oedema | No |  |
| hydrocephaly | No |  |
| Chylous ascites | No |  |
| Spina bifida | No |  |
| **Screening for visible pre-weaning defects** |  |  |
| Body size | Normal |  |
| Skin colour | Normal |  |
| Blotchy coat | No |  |
| Coat colour | Normal | black |
| Belly spot | No |  |
| Head blaze | No |  |
| General activity | Normal |  |
| Tremors /fits | No |  |
| Circling | No |  |
| Head weaving | No |  |
| Ataxia/ gait | No |  |
| hydrocephaly | No |  |
| **Screening for visible weaning defects** |  |  |
| Body size | Normal |  |
| Eye-size/ colour | Normal |  |
| Ear-size/ position | Normal |  |
| Coat colour/texture | Normal |  |
| Skin tension | Normal |  |
| Greasy/rough coat | No |  |
| Curly coat/ whiskers | No |  |
| Thinning/balding coat | No |  |
| Dark footpads | No |  |
| General activity | No |  |
| Tremors/fits | No |  |
| circling | No |  |
| head weaving | no |  |
| Ataxia/gait | Normal |  |
| micrognathia | No |  |
| Short/wide/ thin head | No |  |
| scoliosis | No |  |
| Tail bend | No |  |
| Poly/syndactyly | No |  |
| Fused toes | No |  |
| Limb shape, length | No |  |
| Puffy limbs/ tail | No |  |
| Belly spots | No |  |
| Head blaze | No |  |
| Coat colour | normal | Black |
| hydrocephaly | No |  |
| **5+ weeks** |  |  |
| Breeding | Yes | See GA passport (Supplementary data 4) |
| Litter size | normal | Average 6.8 which is the same as wild type C57/BL6. |
| Litter frequency | normal |  |
| Pre wean loss | 0% |  |
| Diet and preference | N/A |  |
| Gross behavioural defects | No |  |
| Gross cognitive defects | No |  |
| Aggressive behaviours/ group housed | No |  |

Locomotor and metabolic activity assay was measured in using a calorimetric system (LabMaster; TSE Systems; Bad Homburg, Germany). Animals (8 male wild-type mice, 8 female wild-type mice, 8 male Gal5.1 KO and 8 female Gal5.1 KO mice) were placed in a temperature-controlled (20-22°C) room. Animals were placed for adaptation for 1 week before starting the measurements. After calibrating the system with the reference gases (O2, CO2 and N2), the metabolic rate was measured for 7 days and recorded using CalR software(FULL PROTOCOL can be provided on request). No differences in the following measured variables; oxygen consumption, food consumption, water consumption, body temperature and CO_2_ production was observed. Furthermore data was analysed using CALR online software and showed no gross differences for energy expenditure, respiratory quotient (VCO2/VO2), locomotor activity, ambulatory activity or circadian rhythm. Data will not be shown here as an expansion of this analysis will be provided in a future manuscript.

CalR: A Web-based Analysis Tool for Indirect Calorimetry Experiments Amir I Mina, Raymond A LeClair, Katherine B LeClair, David E Cohen, Louise Lantier, Alexander S Banks Cell Metabolism, 2018; doi: http://www.sciencedirect.com/science/article/pii/S1550413118304017

Full cognitive analysis and Clinical biochemistry was not carried out for this study but is planned as part of future experiments.

Taste aversion. Standard (See supplementary data figure 5)
