## Supplementary data 4 for "CRISPR disruption and UK Biobank analysis of a highly conserved polymorphic enhancer suggests a role in male anxiety and ethanol intake"

**Line Holder Dr Alasdair MacKenzie. PPL No P5D8BAA7D**

| **1.** | **Name of strain**  **(technical scientific name and the local in-house name)** | **Scientific: ΔGAL5.1**  **In house: ΔGAL5.1** |
| --- | --- | --- |
| **2.** | **Scientific justification for using this line** | **We have found the enhancer region; GAL5.1, that controls the expression of the GAL gene in the hypothalamus where the products of the GAL gene; the neuropeptide Galanin, controls fat and alcohol intake. We used CRISPR genome editing to delete the GAL5.1 enhancer to produce the ΔGAL5.1 line to show that expression of the GAL gene is significantly reduced. We are now testing the ΔGAL5.1 line to determine whether a lack of the GAL5.1 enhancer influence fat and alcohol intake.** |
| **3.** | **Actual Severity level** | **Mild** |
| **4.** | **Background strain** | **C57/BL6** |
| **5.** | **Originating Establishment/Source** | **Generated in house July 2015** |
| **6.** | **Details of the genetic manipulation** | **CRISPR genome editing using sgRNA guides flanking the GAL5.1 enhancer. Produced by cytoplasmic injection of sgRNA and CAS9 mRNA into one cell mouse embryos which were subsequently oviduct transferred into receptive CD1 mothers.** |
| **7.** | **General Information: coat colour; special housing requirements; single housing aggression; maternal behaviour etc** | **Females group housed and males group housed until after mating when single housed.** |
| **8** | **Notes on enrichment preferences for line.** | **No specific enrichments required compared to WT animals** |
| **9.** | **Phenotypic abnormalities and observable traits with welfare implications i.e. adverse effects; physical abnormalities etc** | **None, possible reduction in high fat food intake.**  Gal(het) x (Gal) het 2018 Total born: 140  None |
| **10.** | **Special considerations for animal health and care including remedial actions** | **Standard precautions as listed in 19b4 and 19b5 of PPL 70/9029;**  **Where the immune status of the animals might compromise health, they will be maintained in a barrier environment. Animals exhibiting any unexpected harmful phenotypes will be killed, or in the case of individual animals of particular scientific interest, advice will be sought from the local Home Office Inspector.**  **When work under terminal anaesthesia is involved, the level of anaesthesia will be maintained at sufficient depth for the animal to feel no pain** |
| **11.** | **Breeding strategy and performance i.e. litter size; litter frequency; pre weaning loss; colony maintenance ie heterozygous, homozygous** | Colony maintained on a mutant homozygous background.  In late 2016 we changed this to setting up as Ko x (OUR HOME BRED C57) to produce heterozygotes. From then onwards we have been setting up as Het x Het as ko and wt mice are needed for experiments/ keeping the colony going in this manner.  As we did not set up Ko x c57 until December 2016 we have no data for 2016.  **Update April 2019 we are now breeding Ko x Ko.**  GAL x GAL Gal X C57(heterozygous colony required) 2015 Total born: 23 2017 Total born: 33 Average litter size: 5 Average litter size: 6.6 Average No,litters/3 months: Average No,litters/3months: Pre wean loss: 0% Pre wean loss:0%  GAL x GAL GAL(het) x GAL(het) 2016 Total born: 91 2017 Total born:215 Average litter size: 6.5 Average litter size: 7.3 Average No,litters/3 months: Average No,litters/3 months:2.5 Pre wean loss :4.3% Pre wean loss : (1 in total)  Gal(het) x (Gal) het 2018 Total born: 140 Average litters size:7.7 Average No of litters per 3 months:4.5 Pre wean loss:1.4% |
| **12.** | **Any additional information e.g. anaesthetic sensitivity** | **None detected** |
