## Supplementary data 5 for "CRISPR disruption and UK Biobank analysis of a highly conserved polymorphic enhancer suggests a role in male anxiety and ethanol intake"

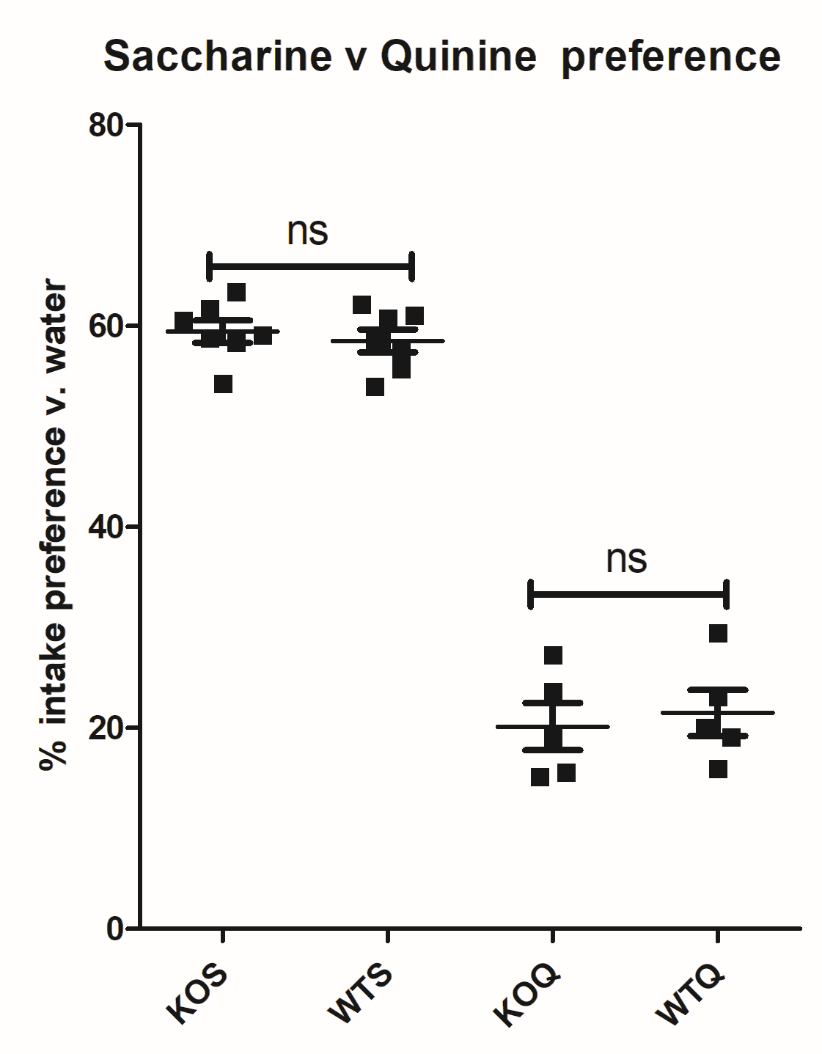

**Supplementary data 5.** Scatterplot demonstrating the percentage fluid intake by singly housed male wild type (WT) or GAL5.1KO (KO) (4-6 month old, *n*=5-7) given a choice of drinking from either a bottle containing a Saccherine solution (KOS, WTS; 2mM solution) or a quinine solution (KOQ, WTQ; 0.3mM) and a bottle containing water. This test was run for 7 days with each bottle being weighed every morning. n.s.=not significant (*p*>0.05). (KOS v WTS; *p*=0.5667, t=0.5891, df=12, KOQ v.WTQ; *p*=0.6898, t=0.4139, df=8)
