## Supplementary data 6 for "CRISPR disruption and UK Biobank analysis of a highly conserved polymorphic enhancer suggests a role in male anxiety and ethanol intake"

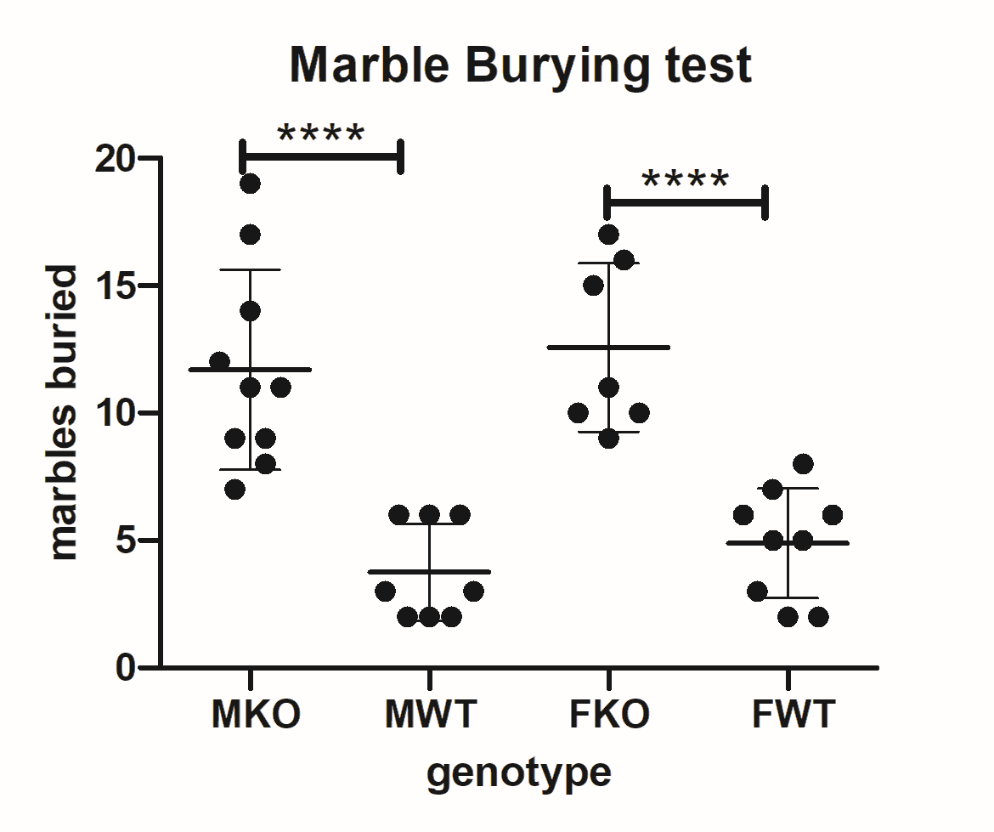

**Supplementary data 6. Marble burying test (MBT).** Scatter plots demonstrating the number of marbles buried in 30 minutes by male knockout (mKO), male wild type (mWT), female knockout (fKO) and female wild type (fWT) mice as analysed by the marble burying test (MBT; n=7-10; F=19.28; d.f.=3). The MBT consisted of a “Eurostandard type iv s” (Techiplast) cage (480 x 375 x 210 mm) cage with 5cm depth of wood chip bedding onto which 20 evenly spaced marbles were placed. Animals were then placed in the cage and the numbers of marbles buried after 30 minutes were recorded.
